# Molecular chaperones inject energy from ATP hydrolysis into the non-equilibrium stabilisation of native proteins

**DOI:** 10.1101/146852

**Authors:** Pierre Goloubinoff, Alberto S. Sassi, Bruno Fauvet, Alessandro Barducci, Paolo De Los Rios

## Abstract

Protein homeostasis, namely the ensemble of cellular mechanisms collectively controlling the activity, stability and conformational states of proteins, depends on energy-consuming processes. *De novo* protein synthesis requires ATP hydrolysis for peptide bond formation. Controlled degradation by the chaperone-gated proteases requires ATP hydrolysis to unfold target proteins and render their peptide bonds accessible to hydrolysis. During and following translation, different classes of molecular chaperones require ATP hydrolysis to control the conformational state of proteins, favor their folding into their active conformation and avoid, under stress, their conversion into potentially harmful aggregates. Furthermore, specific ATP-fueled unfolding chaperones can dynamically revert aggregation itself. We used here various biochemical assays and physical modeling to show that both bacterial chaperones GroEL (HSP60) and DnaK (HSP70) can use the energy liberated by ATP hydrolysis to maintain proteins in their active state even under conditions that *do not favor*, thermodynamically, the native state. The energy from ATP hydrolysis is thus injected by the chaperones in the system and converted into an enhanced, *non-equilibrium steady-state* stabilization of the native state of their substrates. Upon ATP consumption, the chaperone substrates spontaneously revert to their equilibrium non-native state.

## Introduction

Anfinsen *et al*.^1^ demonstrated that artificially unfolded proteins can refold spontaneously into their native, active state without assistance from other macromolecules, solely guided by their primary amino-acid sequences. These seminal *in vitro* refolding experiments implied that, compared to unfolded nascent polypeptides, natively folded proteins are thermodynamically more stable conformations positioned at the bottom of the free energy landscape. This can be reasonably expected from 3.5 billion years of evolution that strived to produce optimally active proteins capable of sustaining long-lasting cellular functions in fluctuating stressful environments. Nonetheless, Anfinsen also noticed that, *in vitro*, the spontaneous folding process can be prone to errors as not all artificially unfolded proteins could efficiently reach their native state, and formed instead inactive insoluble aggregates. Indeed, although the funnel-like structure of the free-energy landscape of proteins should in general favour their native folding^2,3^, local minima, corresponding to metastable misfolded conformations, can transiently halt polypeptides on their way to the native state. There, the kinetically trapped species might pursue further stability by coalescing into stable inactive aggregates, as in the case of alpha-synuclein fibrils and beta-amyloids. Aggregates can be stabilised by intra- and inter-molecular contacts of otherwise inappropriately exposed hydrophobic surfaces and are typically enriched in cross β-sheets^4^, and at sufficiently high protein concentrations their free-energy might be lower than the native state^5−7^. Moreover, mutations and environmental stresses, such as heat-shock, can perturb the free-energy landscape and render the native state intrinsically less stable than other partially compact, misfolded and aggregated conformations^8^. Because protein aggregates and their misfolded precursors appear to be the primary cause for various degenerative diseases, they have gained great interest in the last decades^9,10^.

Molecular chaperones have come to prominence because of their role in assisting *de novo* protein folding and in preventing under stress the formation of toxic aggregates, by several complementary mechanisms involving protein binding and unfolding^5,11,12^. Attesting for their foremost function in sustaining cellular life, up to 3−5% of the total protein mass of unstressed non-cancerous metazoan cells may be constituted from the conserved families of ATPase chaperones, namely the 60 kDa Heat Shock Protein, Hsp60 (GroEL, the bacterial chaperones in parenthesis), Hsp70 (DnaK), Hsp100 (ClpB) and Hsp90 (HtpG)^13^. This amount may further increase following heat stress^14^, or in immortalized cancer cells, which, owing to their abnormal elevated chaperone load, resist apoptosis and protein damage under various chemotherapeutic and physical stresses^15^. Because of their affinity for water-exposed hydrophobic amino acids, chaperones preferentially bind non-native proteins exposing hydrophobic surfaces, thereby passively preventing protein aggregation^16−18^. The role of ATP hydrolysis by chaperones has been traditionally, but possibly over simplistically, associated with the directional cycling through several different molecular conformations, each optimally primed for one of the various steps of the chaperone action: binding, “processing” and release of their to-be refolded, or to-be prevented from aggregating^19^, polypeptide substrates.

Quite surprisingly, a fundamental consequence of the ATPase activity of chaperones, which stems from fundamental laws of physics and remains largely under-appreciated, is that the constant injection of energy from ATP hydrolysis, aside from generating heat, could also drive the system into a *non-equilibrium steady state*, where proteins interacting with the chaperones would distribute over their conformational ensemble differently than at equilibrium^20−23^. A testable prediction in such scenario is that, using the energy of ATP hydrolysis, chaperones should be able to favour the native state of their substrates even under adverse denaturing conditions, where the native state is thermodynamically less stable than the various unfolded, misfolded and aggregated species.

Here, we primarily focussed on bacterial GroEL, which was the first chaperone to be studied *in vivo*^24^ and *in vitro*^16^ and thereafter thoroughly characterized structurally and functionally^25,26^ and on malate dehydrogenase (MDH), which is an extensively studied natural substrate of GroEL in mitochondria and bacteria, as well as on citrate synthase, another well-studied model protein. Further, we show that the same non-equilibrium principles likely apply both to the network of chaperones, composed by GroEL and GroES, the disaggregase ClpB and DnaK, that in cells supervises protein homeostasis, and to Hsp70 (DnaK) alone, which is an abundant ATPase chaperone representing at least 0.5% of the total protein mass of animal cells^13,15^. We find that similarly to GroEL, these systems can also inject free energy from ATP hydrolysis into labile protein substrates and maintain them in a *non-equilibrium* native steady state under denaturing conditions, by mechanisms that, although very different from GroEL at the molecular level, are likely analogous in their overarching principle.

## The free-energy landscape of proteins in non-native conditions

Purified MDH from porcine mitochondria, which is the model protein that we used in most of the experiments, is stable at 25°C, while it spontaneously loses its activity at more elevated temperatures^27^, such as 37°C^28^. These results suggest a picture of the free-energy landscape of MDH at 37°C as represented in Fig.1, where the native, active dimer (ND) is in kinetic equilibrium with the compact, native monomer (NM), which is in turn in kinetic equilibrium with a less-structured intermediate (I) characterized by a higher free-energy. The intermediate is likely short-lived, and it is in kinetic equilibrium also with a collapsed but non-native misfolded state (M)^29−31^, which is more stable *(i.e.* lower in free-energy) even than the native dimer.

**Figure 1.**
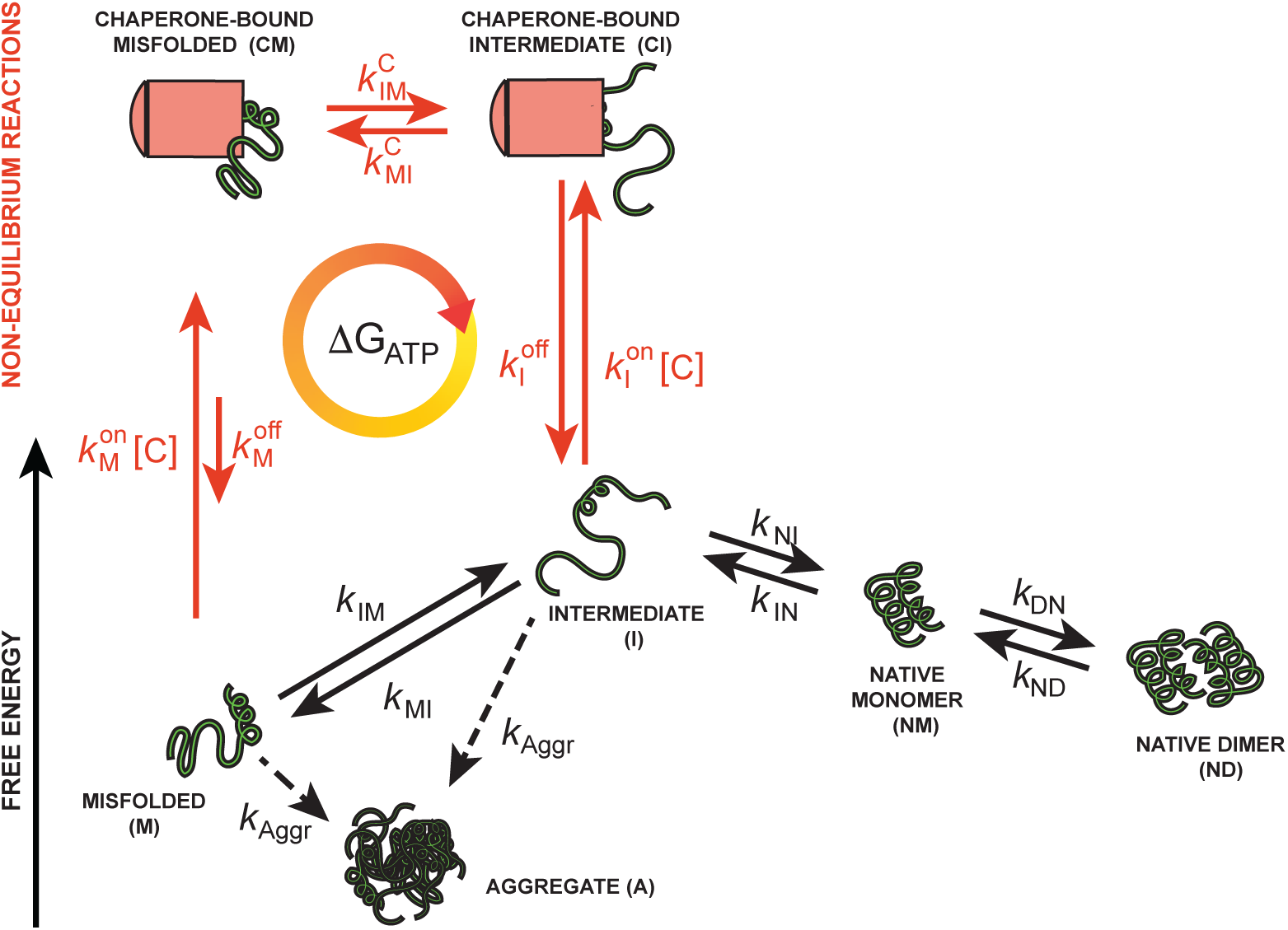
Free energy landscape of MDH at 37°C and schematic action of chaperones. The free-energy landscape of MDH at 37°C is represented by the native (active) dimer (ND), the native (inactive) monomer (NM), a less structured intermediate (I) which is higher in free energy and a misfolded state (M), which is the lowest in free-energy. Non-native conformations can further lead to the irreversible formation of aggregates (A), although likely at much slower rates. The spontaneous conversion reactions between the different states are represented with black arrows (dashed arrows for aggregation to highlight the different timescale of this process). Chaperones act on the non-native ensemble by associating to and dissociating from misfolded (CM) and intermediate (CI) conformations. While bound to chaperones, misfolded and intermediate conformations can interconvert into each other. Reactions driven by chaperones are represented by red arrows. The energy driving the chaperone action is graphically indicated with the yellow-to-red closed-circle arrow.

Aggregation, which is a concentration dependent reaction, further proceeds by the binding of non-native monomers to other non-native monomers or to already formed aggregates (A). Since it has been noted that non-native MDH remains in conformations that, when brought back into native conditions, could refold spontaneously in native conditions even after incubation at denaturing temperatures for tens of minutes^32^, the aggregation is very likely a slow process compared to denaturation at micro-molar, or lower, concentrations. These findings also agree with observations that several non-native proteins can remain soluble for long periods of time, even under adverse conditions that do not favour the native state^8,33,34^.

## Under denaturing conditions GroEL and ATP maintain MDH in a *non-equilibrium* native state

Hartman *et al*.^28^ remarkably found that at 37°C the rate of spontaneous denaturation of MDH decreased in the presence of substoichiometric amount of GroEL and GroES 7-mers and ATP. These intriguing observations did not trigger a more fundamental issue: does GroELS use ATP hydrolysis to seemingly “preserve”, under a denaturing temperature, a conformational state that is not the most stable one as dictated by thermodynamic equilibrium? We address here this crucial question, which has remained to date unanswered.

At 25°C, as expected, native MDH remained stably active with or without GroELS and ATP (Fig.S1A). At 37°C, MDH spontaneously lost its active form at an approximate rate of 0.007 min^-1^ (Fig.2A), compatibly with Hartman *et al*.^28^ and with the scheme in Fig.1. Addition of a molar excess of GroELS (3.5 *µ*M protomers of GroEL and GroES) (but without ATP) over MDH did not affect the rate of denaturation (Fig.S1B). In contrast, when GroELS, ATP and a pyruvate kinase/phosphoenolpyruvate-based regeneration system were supplemented at t=0’, they caused a net reactivation of nearly 75 nM MDH on top of the initial amount of active enzyme. This implied that in the initial stock of nominally native MDH there was at most 80–85% of truly native, active MDH species, alongside ~15–20% inactive yet soluble and GroELS-amenable species, in keeping with the results of Peralta *et al*.^3^*^2^*, since no protein aggregate has yet been shown to be solubilized by GroELS+ATP^12,35^.

**Figure 2.**
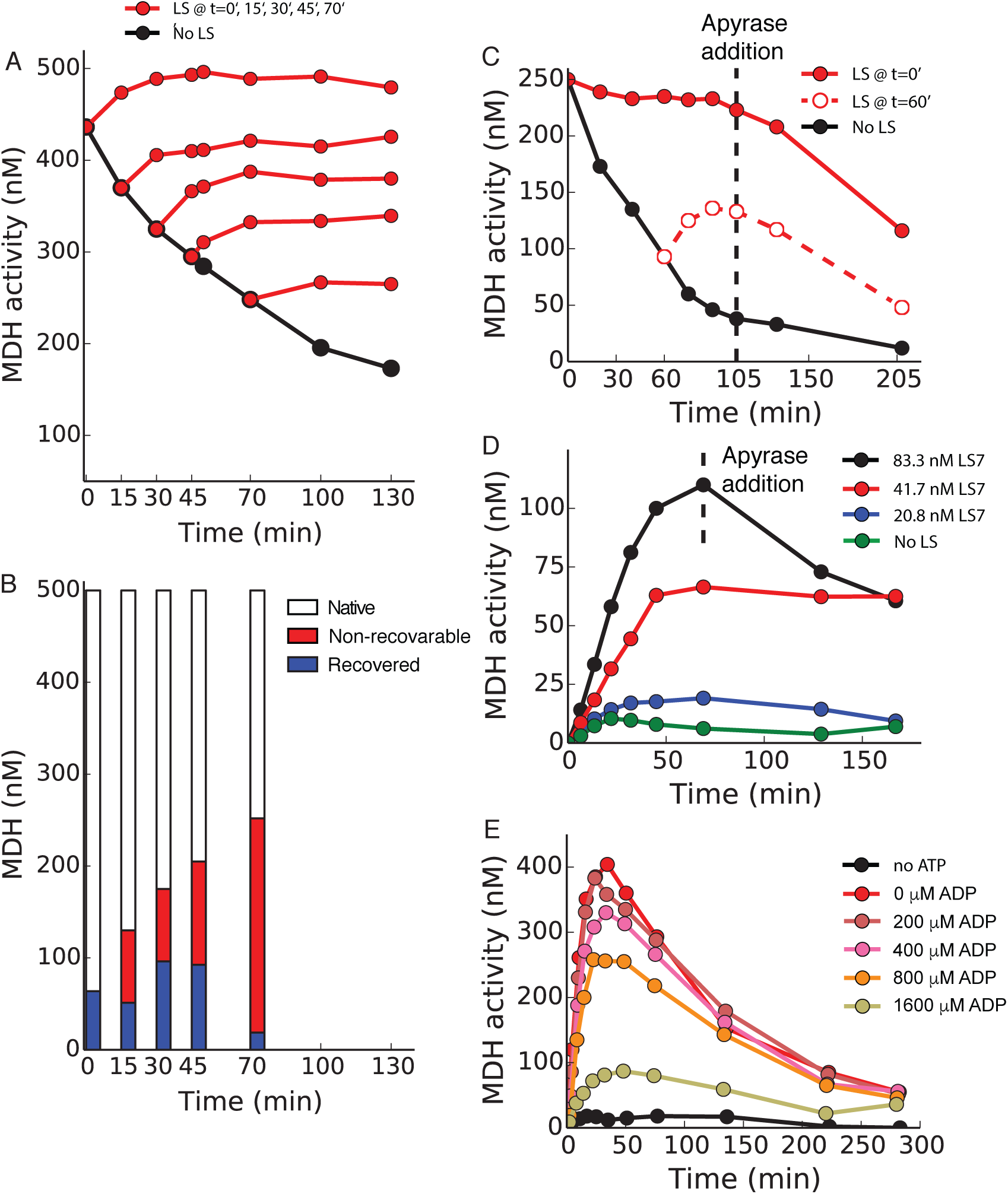
GroELS recovers and then maintain MDH activity in denaturing conditions, in a stringently ATP-dependent, non-equilibrium, reaction. A) Time course of the activity of MDH (0.5 µM) at 37°C in the absence (black circles) and presence (red circles) of GroEL (3.5 µM) and GroES (3.5 µM) and ATP regeneration system (see Methods). In the absence of GroELS MDH spontaneously lost activity at an apparent rate of about 0.007 min^-1^. GroELS was supplemented at different t=0’, 15’, 30’, 45’ and 70’ (see Methods for the choice of the scale on the y-axis; the same procedure applies to all graphs where activity is expressed as a concentration). B) Final MDH fractions (at 130’) for GroELS addition at different times from the experiments in Fig.1A. White bars represent the concentration of active MDH at the time of GroELS addition, whereas the recovered fractions (blue bars) are the differences between the active protein concentrations at 130’ and at the time of GroELS addition. The concentration of proteins that cannot be recovered by GroELS is represented with red bars. C) GroELS (3.5 µM) and ATP regeneration system were added to 0.25 µM MDH at 37°C (denaturing conditions) at t=0’ (red filled circles) or at t=60’ (red empty circles). Apyrase was added at t=105’. D) Refolding of urea-pre-denatured 0.5 µM MDH at 37°C in the initial presence of different GroELS concentrations (as indicated) and ATP regeneration system. Apyrase was added at t=60’. E) Refolding of urea-pre-denatured 0.5 µM MDH at 37°C in the initial presence of GroELS (3.5 µM), 400 µM ATP and various concentrations of ADP (as indicated). As predicted by the correct accounting of the energy available for the chaperone action (see main text), increasing concentrations of ADP progressively inhibit the reaction.

Surprisingly, following reactivation MDH remained active and apparently stable, despite the denaturing conditions in which the active state should not have persisted. When GroELS and ATP were added later after the start of thermal denaturation, native state maintenance again followed the net reactivation of some MDH (Fig.2B), which had remained chaperone-amenable during 30–60 minutes of the heat treatment. The amounts of recovered MDH significantly decreased only when GroELS and ATP were added at much later times, once more suggesting that the formation of aggregates (Fig.2B, non-recoverable fraction) was much slower than the denaturation process in our assay conditions.

Our observations highlighted that at 37°C, where the non-native ensemble is more stable than the native ensemble (Fig.1), the primary ATP-dependent action of GroELS was not limited to the mere prevention of aggregation of the non-native MDH. Instead, as represented in Fig.1, GroELS behaved as if it actively rescued the recoverable misfolded MDH conformers, pushing them up the free-energy landscape towards the native active state, where, despite being only metastable, they were maintained for extended periods. These conclusions were further strengthened by a similar action of GroELS+ATP during heat denaturation of citrate synthase, another classical GroEL substrate (Fig.S2)^36^.

Basic physics principles dictate that the steady maintenance of MDH in its otherwise unstable native state must be strictly ATP-dependent. Indeed, if no energy could be extracted from ATP hydrolysis, equilibrium thermodynamics would inescapably govern the system, and GroELS could not oppose the thermodynamically inevitable denaturation of the MDH. We thus devised experimental setups where we could control whether the system was at, or away from, equilibrium.

As in Fig.2A, an excess of GroELS over MDH, supplemented with ATP and a regeneration system, was expectedly able to maintain the native MDH level when added from the start and, when added at a later stage, to recover some activity and then maintain it constant despite the denaturing temperature (Fig.2C). In both cases, the subsequent addition of apyrase, which readily hydrolysed ATP into AMP, deprived GroELS of its energy source, inhibiting its ability to maintain the non-equilibrium active stabilisation of the metastable native MDH at 37°C. As a consequence, the system spontaneously sought the MDH inactive denatured state, which is thermodynamically the most stable (by contrast, at 25°C, where native MDH is intrinsically stable, addition of apyrase had no effect, see Fig.S1A).

Using MDH that was first artificially unfolded by urea, Anfinsen-type *in vitro* refolding experiments showed that, following removal of urea by abrupt dilution, there was no spontaneous MDH refolding at 37°C (Fig.2D), consistent with the thermodynamic instability of the native state at that temperature. In contrast, we found that the presence of GroELS+ATP at the time of the urea dilution resulted in the transient major accumulation of native MDH at 37°C, and in its subsequent maintenance over time. Apyrase addition readily brought the level of native MDH back under the control of equilibrium thermodynamics, and led to its spontaneous denaturation despite the presence of the chaperones (Fig.2D).

The amount of energy that is liberated by ATP hydrolysis, ΔG_ATP_, and that is available for GroELS to inject into maintaining the metastable MDH into its native state, depends on the concentrations of ATP, ADP and inorganic phosphate Pi through the relation:

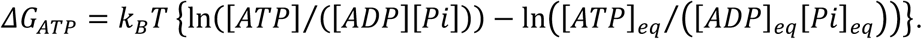

Here, *k_B_* is the Boltzmann constant, *T* is the absolute temperature and the *eq* subscript marks the concentrations that the different molecules would have if the hydrolysis reaction could run to completion (*i.e.* [ADP]_eq_/[ATP]_eq_ ≈ 10^6^™10^8^ depending on the solution conditions^37^). To analyse the role of ΔG_ATP_ in more detail, we changed the concentrations of ADP in the GroELS-mediated refolding experiments of urea pre-unfolded MDH at 37°, while keeping ATP constant at a sub-physiological concentration (400 µM), which was, however, still in excess over GroELS (3.5 µM). Increasing amounts of ADP, from 0 µM to 1600 µM, were added at the start of the reaction, with the goal of progressively reducing the value of ΔG_ATP_, as predicted from the above formula. Expectedly, larger ADP concentrations inhibited the initial effectiveness of the chaperone mediated refolding reaction of the MDH (Fig.2E). Later, the reaction was further hindered by the decreased concentrations of ATP that was consumed, and by the reciprocal increased concentration of ADP. This brought the system back towards equilibrium conditions, namely ΔG_ATP_=0.

Together, the results in Figs.2C–E confirmed that the effect of GroELS is strictly dependent, and tuneable, by the amount of energy available from ATP hydrolysis, which thus drove and maintained the system in a non-equilibrium state.

## Non-equilibrium thermodynamics model of chaperone action

We next rationalized these observations with a model that captured the main properties of the free-energy landscape of MDH and of the ATP-fuelled chaperone action as represented in Fig.1. We chose to describe the action of chaperones at a minimalistic level of detail to focus on the broader and general thermodynamic implications of the energy consumed during their cycle. The explicit reactions of chaperone-substrate association and dissociation allowed accounting for the chaperone concentrations that, when needed, we chose to vary in order to serve our experimental demonstration. When bound to chaperones, substrates could further convert between more and less structured non-native conformations, based on the compelling experimental evidence for Hsp60s (and for Hsp70s and Hsp100s) that following binding, misfolded substrates generally undergo a loss of secondary structure (unfolding) and an expansion, while still in complex with the chaperones^34,38–46^, as also recently emphasized by Dill *et al*.^47^. This assumption was further strengthened by observing that the compacting action of a crowding agent (15% PEG 6000) ^48,49^, which inhibited both the misfolded-to-native and native-to-misfolded conversions of MDH at 25°C and at 37°C (Fig.S5A), was antagonized by GroELS+ATP at 37°C (Fig.S5B). In other words, the energy of ATP hydrolysis was used by GroELS to facilitate the de-compaction and/or unfolding of its substrates.

In this model, the native species do not interact with the chaperones, in agreement with the generally-accepted view that in order to act efficiently in the crowded environment of cells, molecular chaperones must have a high affinity for non-native, aggregation-prone *de novo* synthesized, *de novo* translocated or stress misfolded polypeptides, while keeping an extremely low affinity for the very dense population of surrounding proteins that are native^50^.

The detailed internal movements GroEL, as well as GroES binding and release, the precise steps of ATP hydrolysis and nucleotide exchange, and their cooperative and/or anti-cooperative nature are usually taken into account when addressing the precise molecular steps of the cycle of the action of GroEL^51−54^. Here, they needed not to be explicitly represented in the model, as they were not necessary to extract the effects of the energy consumed by the chaperones on the behaviour of the system. Indeed, according to the laws of non-equilibrium thermodynamics, which must be valid at any coarse-grained level of description, the effect of the injected energy from ATP hydrolysis, ΔG_ATP_, could be rigorously integrated in the relations between the association and dissociation rates of the chaperone-substrate complexes (see SI).

The rates of the transitions between the different MDH states, the rate of aggregation and the rates of the chaperone action have been chosen to reasonably reproduce the measured spontaneous denaturation of native MDH dimers at 37°C and the third reactivation curve (when GroELS and ATP were added at t=30’) observed in Fig.2A (see SI for the full list of rates). All model results presented in this work have been obtained with this same set of intrinsic parameters.

Once the parameters had been set, we found that the model closely reproduced the other data in Fig.2A, when the GroELS was added either earlier or later during the thermal denaturation *(cf.* Fig.3A). More importantly, the model gave access to estimates of the various populations of MDH species that were not directly detectable experimentally (Fig.3B). The chaperone-resistant non-recoverable fraction was heterogeneous: it comprised an irreversibly inactive component, likely aggregates, which expectedly grew over time at 37°C, and a proportionally diminished soluble component, which was mostly composed of near-native inactive monomers.

**Figure 3.**
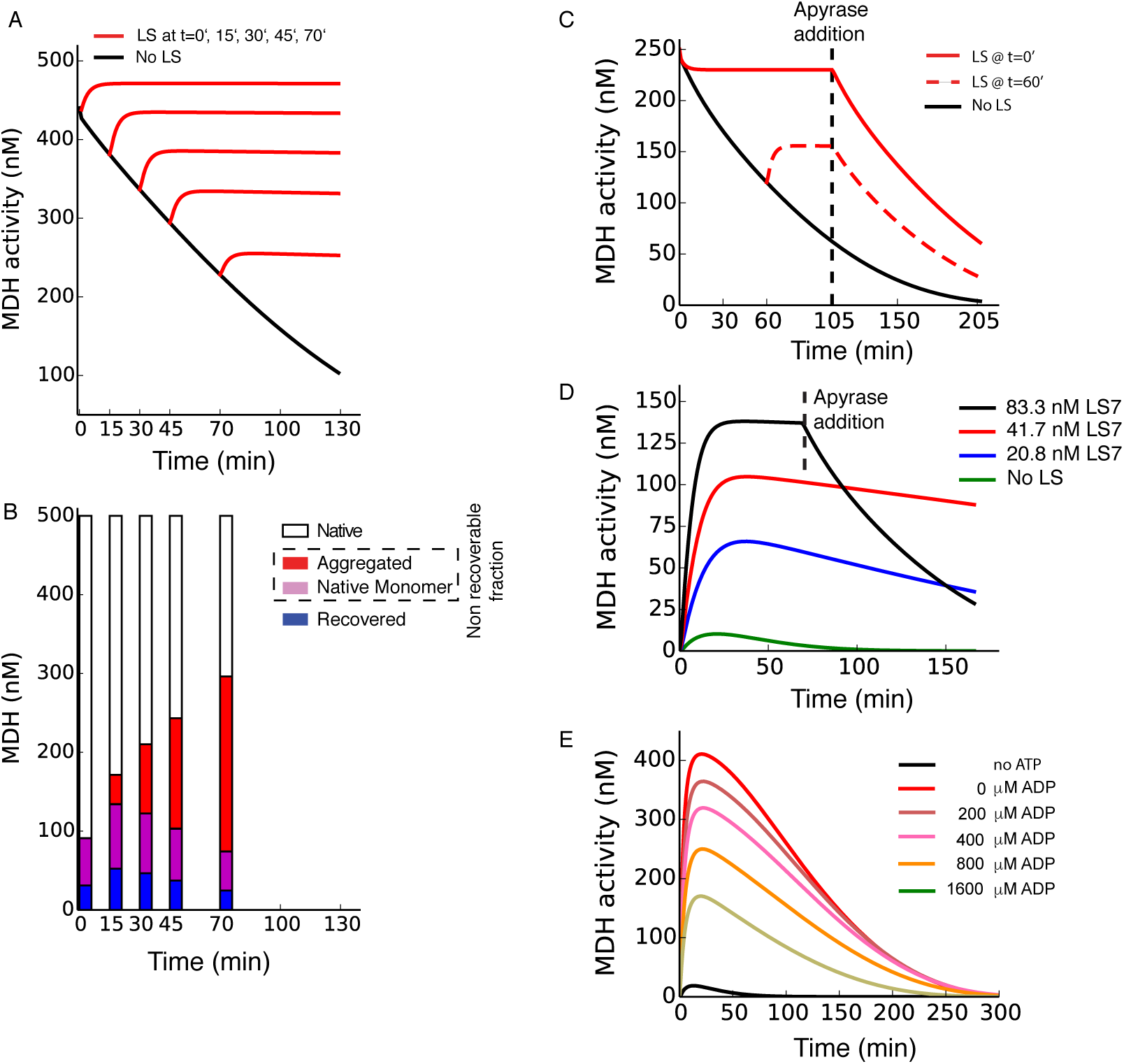
Results from the model. The panels here are in one-to-one correspondence to the panels in Fig.2, and have been obtained for a single set of intrinsic parameters (the rates of the reactions). Concentrations of chaperones and substrates are the same as in the corresponding experiments. A) and B) Model results for the same setups as in Figs.2A and 2B respectively. C) Results from the model for the same conditions as in Fig.1C. D) Results from the model for the same conditions as in Fig.2D. E) Results from the model for the same conditions as in Fig.2E (see SI for details on the modelling of ATP consumption).

In the model, the switch from non-equilibrium to equilibrium conditions induced by apyrase could be captured by sharply setting ΔG_ATP_=0 at the time of apyrase addition, and the results agreed with the experiments (compare Fig.3C with Fig.2C). Similarly, we could reproduce the results from Anfinsen-type refolding of artificially unfolded MDH experiments (Fig.3D). In the model, we could also control the amount of energy available to chaperones by changing the [ATP]/[ADP] ratio and taking into account its variation in time (see SI for the implementation in the model). The same behaviour observed experimentally in Fig.2E. was reproduced by the model (Fig.3E), further confirming that the energy of ATP was transduced into the stability of the native state of MDH.

## GroELS acts through iterative cycles fuelled by ATP hydrolysis

We then investigated whether the observed ATP-dependent ability of GroELS to regenerate and transiently maintain native MDH under denaturing conditions (Fig.1A) was possibly due to the stabilisation during long-lived in-cage binding, of a minority of denaturing polypeptide substrates by a molar excess of GroELS.

We thus exposed during 8 hours, an MDH concentration higher than in previous experiments (2.5 µM) to decreasing substoichiometric amounts of GroELS in the presence of a non-limiting amount of ATP (backed by a long-lasting ATP regeneration system) (Fig.4A). Following 8 hours at 25°C, the native MDH expectedly remained nearly fully native, independently of the presence or absence of GroELS. Without ATP, all GroELS concentrations remained ineffective at preserving at 37°C the MDH activity and 85% of the total MDH, *i.e.* about 2.1 µM, became inactive in 8 hours. However, in the presence of ATP, as little as 357 nM of GroEL_7_ rings (2.5 µM of GroEL protomers) could actively maintain a 7-fold excess of MDH in the native state during the 8 hours at 37°C. At EC_50_, as little as 56 nM of GroEL_7_ rings (392 nM of GroEL protomers) could effectively maintain in the native state as many as 923 nM of MDH, implying that on average, each GroEL7 cavity had actively and iteratively converted at least 17 molecules of inactive MDH monomers into active dimers, more than it could bind at any given time during the 8 hours (Fig.4A, inset). This ruled out the possibility that the stabilisation of the native species resulted from the lengthy association with the chaperone cage. Without changing parameters, the model could closely reproduce these experimental data (Fig.4B), including the minimal number of recovered MDH molecules, which was about 20 per GroEL_7_ ring at EC_50_ (Fig.4B inset, red squares). Moreover, because at any moment of the 8 hours reaction each MDH monomer that was released from the chaperone cavity and that had natively refolded could denature again, the model further unravelled that each GroEL_7_ cavity had to undergo at least 400 ATP-fuelled cycles of MDH binding, processing and release in order to yield the measured elevated end-product amounts of native MDH after 8 hours at 37°C (Fig.4B inset, blue squares).

**Figure 4.**
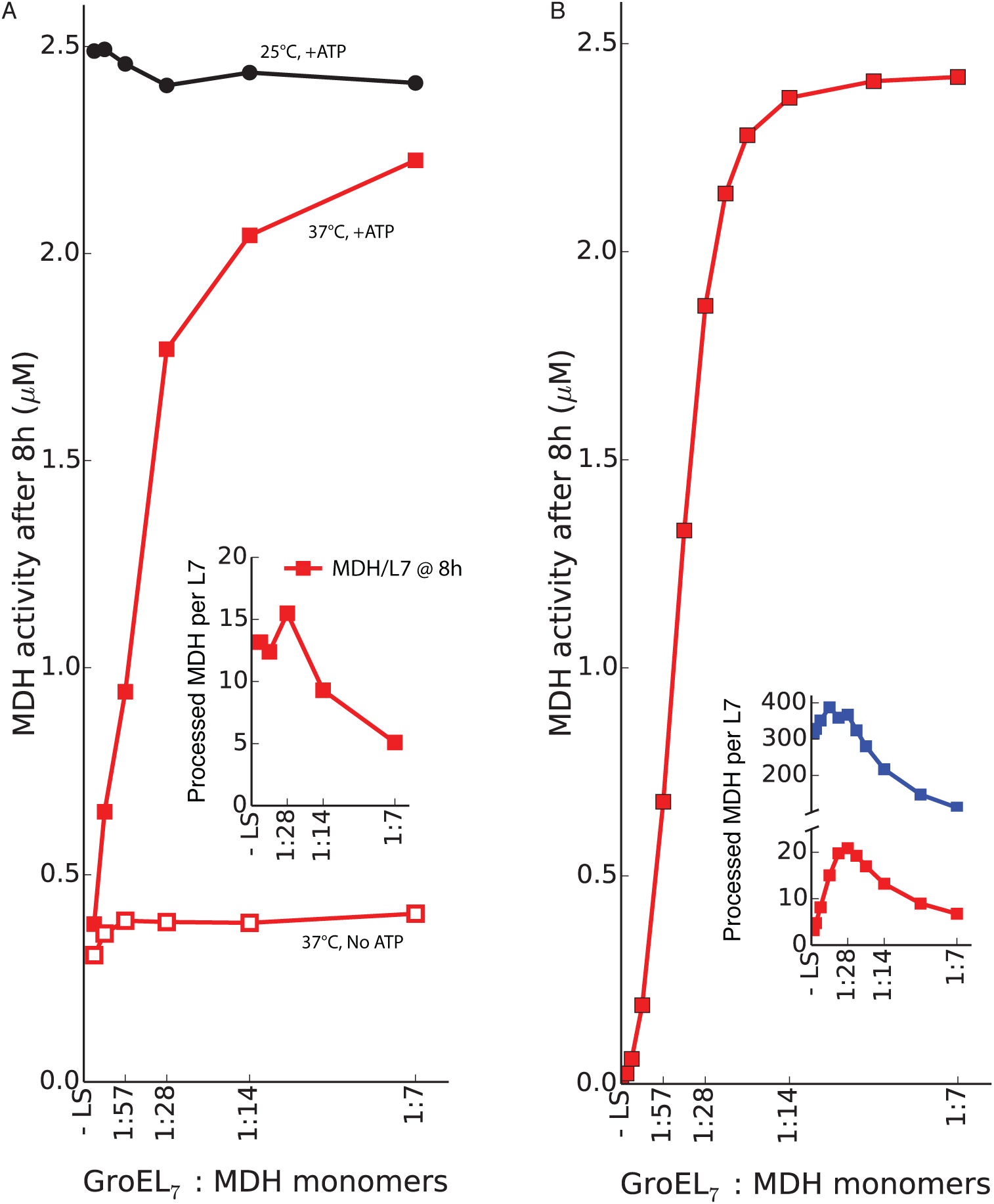
ATP-driven catalytic action of substoichiometric amounts of GroELS on MDH at 37°C. A) Activity of 2.5 µM MDH after incubation for 8 hours in the presence of increasing amounts of GroEL (and equimolar GroES) at 25°C (black circles) and at 37°C in the presence (full red squares) and absence (empty red squares) of ATP and regeneration system. The ratio of the concentration of GroEL_7_ rings (considered as the minimal active unit of GroEL) to the concentration of MDH protomers is given. Inset) The minimal number of non-native MDH protomers that should be rescued by each GroEL_7_ ring to explain the observed activity after 8h. B) results of the model in the same conditions as in panel A. Inset) The minimal number of non-native MDH protomers that should be rescued by each GroEL_7_ ring in order to explain the observed activity after 8h (red squares) is similar to the experimental one (~15 protomers per GroEL_7_). The model gives also access to the effective number of MDH protomers that had to be processed by each ring (blue squares), which is more than one order of magnitude larger (~300 per GroEL_7_ at EC50), highlighting the iterative, catalytic action of GroELS on its substrates.

These results imply that continuously misfolding polypeptides must have been effectively converted into native complexes against thermodynamic equilibrium through a highly iterative process which is similar to catalysis, because the conversion between the misfolded and the intermediate state is much faster than it would be spontaneously, and where the catalyst molecule (here, GroELS) is available for multiple turnovers of the reaction. Yet, at variance with catalysis, this process is not accelerated in both directions because of energy consumption, leading to the observed change of the native state fraction in steady-state.

## GroELS uses the energy of ATP hydrolysis to shape the effective free-energy landscape of proteins

The results in Figs.2 and 4 indicate that by readily processing the thermally misfolded proteins as they formed and by releasing the processed species to transiently form native complexes, GroELS and ATP were in fact maintaining the concentration of the aggregation-prone misfolded proteins very low, and consequently dramatically reduced the process of aggregation that strongly depends on the concentration of misfolded precursors. This justified our further simplification of the model by plainly neglecting the contribution of aggregation. It then became possible to formally define the steady-state *effective* free-energy difference between the native and non-native ensembles, 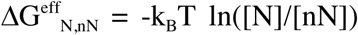 (see SI for a discussion) as a function of ΔG_ATP_. Here, [N] was the total concentration of native conformers ([N]=[NM]+[ND]) and [nN] the total concentration of non-native conformers ([nN]=[I]+[M]+[CI]+[CM]) (see Fig.1). Expectedly, at 37°C, in equilibrium conditions (ΔG_ATP_=0) the non-native ensemble was more stable than the native ensemble 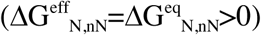, driving the steady spontaneous denaturation of native MDH (Fig.5A). As ΔG_ATP_ increased by injecting progressively more energy in the reaction, the native ensemble became stabilized. For ΔG_ATP_> 2.7 kcal/mol, the native ensemble became more stable than the non-native ensemble. For ΔG_ATP_=7.7 kcal/mol (the value used in the model in Figs.2, 3A–D and 5C), the native-ensemble was found to be apparently more stable than the non-native ensemble by about 1.8 kcal/mol, corresponding to a total energy transfer by the chaperone of about 2 kcal/mol from ATP hydrolysis to the steady-state stability of the native state product. Pictorially, the effect of the action of GroELS thus corresponded to a remodelling of the free-energy landscape of the substrate, where the misfolded state was destabilised and raised above the native state, as it would be expected under physiological conditions (Fig.5B).

**Figure 5.**
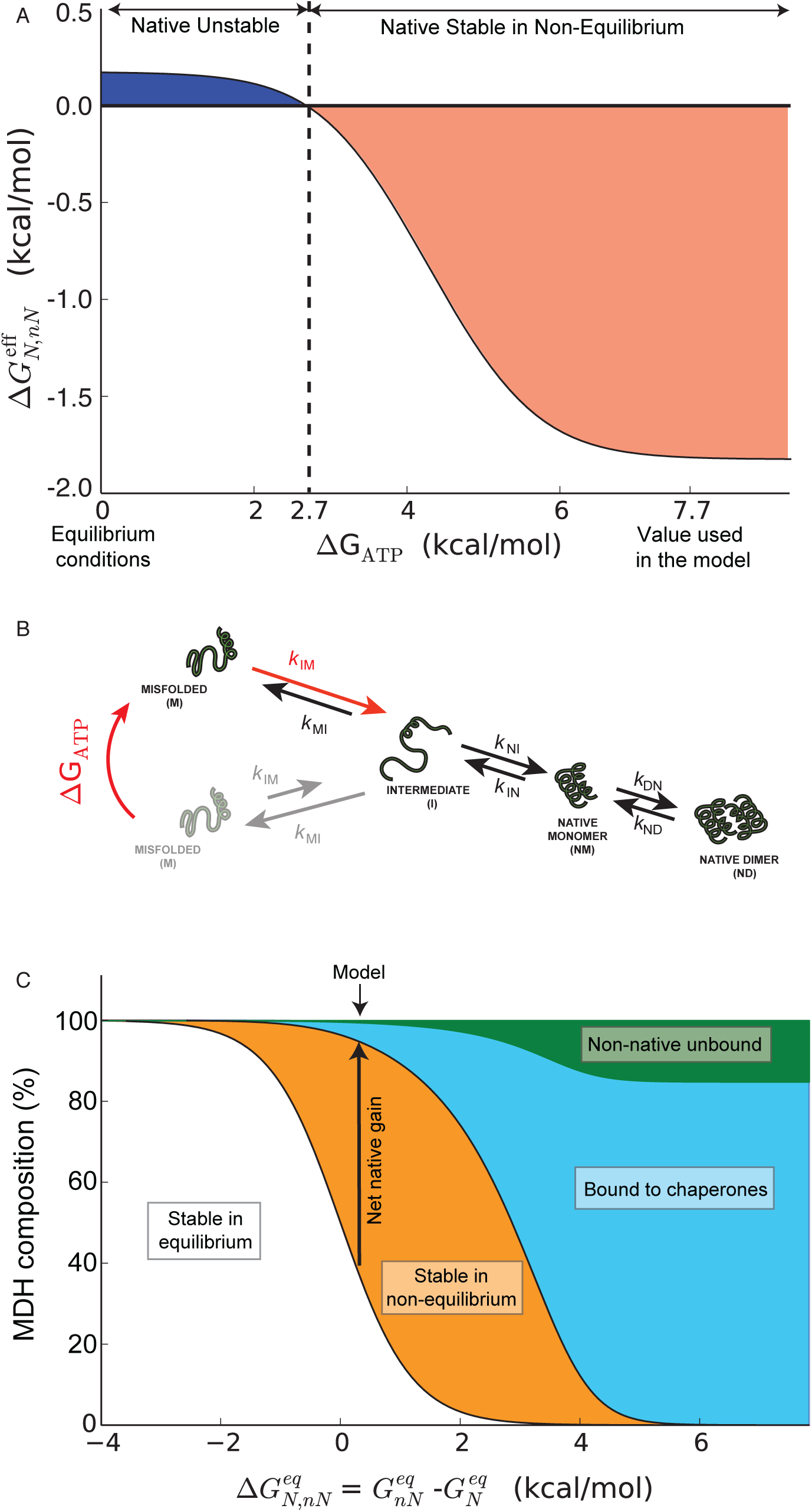
GroELS turn the energy of ATP into a non-equilibrium stabilisation of the native state of MDH. The *effective* free-energy difference between the native and non-native two ensembles 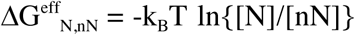 (as defined in the main text) as a function of ΔG_ATP_, from equilibrium conditions (ΔG_ATP_=0) to conditions exceeding the ones that, in the model, reproduce the experiments. 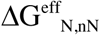 is positive at equilibrium at 37°C, attesting for the thermodynamic instability of the native ensemble. The native ensemble of MDH is stabilized as progressively more energy is injected into the system. B) The action of GroELS on the non-native ensemble in non-equilibrium results can be visualized as a modification of the free-energy landscape of non-native conformations, whereby misfolded proteins have a much higher free-energy than native conformations. C) The relevance of the action of GroELS as a function of the equilibrium stability of their substrates 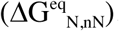. When 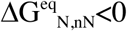, substrates benefit very little from the help of GroELS because they are already stable. As the equilibrium stability of the native state is challenged (increasing 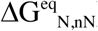), energy injection in the system helps maintaining the native state populated against its thermodynamic tendency (the orange region represents the net gain of native proteins thanks to chaperones). Extreme instability resulting from strong stresses or highly deleterious mutations cannot be compensated by chaperones (blue region), which can anyway avoid protein aggregation by binding the non-native conformations. Non-native but unbound, and thus aggregation-prone conformations thus represent a minor part of the solution (green region).

The model also rationalized the role that an elevated cellular concentration of GroELS (an estimation from quantitative proteomics of a rat liver cells indicates that in the mitochondrial stroma there are about 130 µM HSPD1 (GroEL) protomers, as many HSPE1 (GroES) protomers, for about ~160 µM of MDH2 protomers^13^), might have for the evolution of proteins carrying cellular functions necessitating high flexibility at an unavoidable cost of decreased overall intrinsic stability (Fig. 5C). As shown by Anfinsen, intrinsically stable native proteins 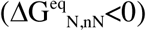 may not need assistance from other proteins. Here, we found, however, that as the thermodynamic stability of labile proteins was progressively challenged by mild stress or mutations (corresponding to increasing values of 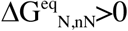), molecular chaperones could recruit the energy of ATP hydrolysis to maintain in their native state, proteins that are intrinsically metastable, thereby extending their effective range of activity within stressed cells (the orange region in Fig.5C represents the net gain of native proteins over the amount that should be present according to equilibrium thermodynamics). Noticeably, even under harsh denaturing conditions, where an excess of GroELS is unable to maintain MDH in the native state, in the model, it was still able to bind the non-native aggregation-prone species (blue region). When GroELS was in excess over the substrate, it could reduce the amount of free non-native species (green region), thereby impeding their irreversible aggregation and justifying, in part, the somewhat overstated characterizing of molecular chaperones as being able to prevent the aggregation of other proteins^55^.

## The full power of energy transduction is unleashed by the entire chaperone network

In bacterial cells, protein homeostasis depends on several ATPase chaperones, comprising GroELS, DnaK (and its co-chaperones), HtpG and the disaggregase ClpB^56^. We thus tested the effects of a more physiological multi-chaperone composition, for various concentrations of DnaK, on MDH at 37°C (Fig.S4). At 37°C, when urea-unfolded MDH was pre-incubated for 70 minutes with ATP, ClpB, GroELS, and the DnaK co-chaperones DnaJ and GrpE, but without DnaK, no significant native MDH refolding was observed. The subsequent addition of increasing amounts of DnaK caused the increasing accumulation of native MDH, despite the non-native temperature, demonstrating that DnaK, which can act both upstream and downstream of ClpB^57^ and of GroELS^58,59^, can facilitate the injection the energy of ATP hydrolysis into increased stability of labile native proteins.

As matter of fact, early *in vitro* experiments with DnaK already indicated that also these chaperones are able to convert the energy of ATP into a non-equilibrium native state stabilisation of the thermolabile luciferase at denaturing temperature^60^. We further demonstrated here that this is an intrinsically non-equilibrium effect, by showing that upon the addition of apyrase, spontaneous denaturation of luciferase was restored (Fig.S3A). We further tested the ability of DnaK to stabilize native MDH at 37°C, and observed that the addition of KJE and ATP produced a significant, although only short-lived, reactivation of native MDH (Fig.S3B). We conclude thus the KJE+ATP chaperone system is also able to transduce the energy of ATP into the non-equilibrium stabilisation of the native state of both Luciferase and MDH at 37°C, even if, in the case of MDH, less efficiently than GroELS.

Together, these results suggest that distinct ATPase chaperones may act individually as well as cooperatively at the effective injection of energy from ATP hydrolysis to artificially maintain labile proteins in their native states under stressful conditions where they spontaneously tend to misfold and aggregate, likely because aggregates represent the true free-energy minimum and are thus thermodynamically most stable^5,6^. In the cell, the conformational landscape could thus be modified even more dramatically than by GroELS alone, by raising the apparent free-energy, not only of misfolded monomeric conformers, as in Fig.5B, but also of already formed stable aggregates, thus further extending the conditions in which labile proteins can remain mostly native.

## Conclusions

Our parallel experimental and theoretical findings have revealed that, at its most fundamental level, the role of ATPase chaperones such as GroEL, DnaK and ClpB, is to transduce the energy liberated by ATP hydrolysis into an enhanced *non-equilibrium* steady-state stability of otherwise labile native protein substrates, even under conditions where their native states are unstable. In this respect, the function of chaperones is more deeply rooted in non-equilibrium thermodynamics than in the molecular aspects of the ATP-driven conformational transitions in the chaperone complexes, that we have neglected here in favour of a simplified model in which we considered only the constraints imposed by thermodynamics, that mandatorily apply independently of the level of simplification of the chaperone cycle. Thus, at the price of missing part of the molecular accuracy characterizing more complex simulation schemes^61,62^, our simpler strategy provided a general rigorous thermodynamic perspective that broadly applies to many ATPase chaperones.

In a cellular perspective, we may thus propose that ATPase chaperones in general supply cells with an energy consuming mechanism to withstand prolonged destabilizing stresses, such as the diurnal cycle of temperatures, by maintaining the most labile members of the proteome in a soluble functional state until the environmental stress is over. Chaperones might thus increase the freedom of evolution to accumulate functionally favourable but destabilizing mutations^63^, by artificially maintaining mutated proteins active and functional in the cell, against their thermodynamic tendency to spontaneously denature. This view is compatible with directed evolution experiments that have demonstrated that, when chaperones are over-expressed, enzymes can evolve faster, remain active and soluble even when challenged by slightly destabilizing mutations^64−67^.

Our observation with labile proteins, together with the emerging evidence that ATP-dependent mechanisms can also actively maintain RNA molecules in steady-state conformations that are different from the ones they seek to adopt at equilibrium^68−70^, lends further support to the general concept of the cytoplasm being an “active milieu”^71,72^. Indeed, when ATP becomes limiting, essential biological functions, such as the maintaining of steep ion gradients across membranes, collapse and lead to cell death. Thus, energy-consuming nanomachines need to be constantly at work to maintain cells in a metabolically active state. In particular, stress-labile macromolecules, such as DNA, RNA and proteins damaged by exposures to various physical and chemical stresses, need to be constantly proofread by enzymes, such as topoisomerases, helicases, RNA chaperones and now, as we show, also by protein chaperones, which, at an energy cost, can repair and convert them to their active, functional conformations. These ATP-fuelled enzymes can thus remodel the free-energy landscape of biological macromolecules, which consequently become less limited by the dictates of equilibrium thermodynamics. This calls for questioning the correlation between the behaviour of marginally stable biological macromolecules observed *in vitro*, as compared to their effective behaviour in the highly crowed, chaperone-rich environment of the cell. From the perspective of the landscape theory of protein folding, which posits that the native state sits at the bottom of a funnel-shaped free-energy basin^2,3^, our results suggest that the presence of other minima, possibly lower in energy than the native state, is not going to affect correct the folding as long as the corresponding conformations can be processed by chaperones.

The present results are part of a growing recognition that all aspects of life, from its origins billions of years ago to its present most intimate workings, depend on a constant influx and subsequent dissipation of energy. Only the accounting of this energy budget, and of the way it is used to offset very complex biological systems away from their natural tendency to reach thermodynamic equilibrium, can lead to a correct quantitative description of biological systems. As Erwin Schrödinger stated, “Living matter evades the decay to equilibrium”^73^. Chaperones represent a vivid incarnation of Schrödinger’s fundamental insight.

## Materials and Methods

### Proteins

GroEL, GroES were purified according to Torok *et al.*^74^. DnaK, DnaJ were purified as according to Gur *et al.*^75^. GrpE was a gift from H.-J. Schönfeld, F. Hoffmann-La Roche, Basel, Switzerland. ClpB was purified according to Woo *et al*.^76^. Firefly luciferase (Luc) was purified and assayed in as described in Sharma *et al*.^42^. Mitochondrial malate dehydrogenase (MDH) was from pig heart mitochondria (Boehringer Mannheim) and assayed according to Diamant *et al*.^77^. Citrate synthase (CS) was from Sigma and assayed according to Buchner *et al*.^36^.

### Chaperone assays

The chaperone-refolding buffer was 100 mM Tris 7.5, 150 mM KCl, 20 mM MgCl2, 10 mM Dithiothreitol. Unless mentioned otherwise, all chaperone assays used 4 mM ATP 5 mM 2-phosphoenolpyruvate and pyruvate kinase (Sigma) to regenerate the ATP.

Concentrations of chaperones, cochaperones and reporter protein: Unless mentioned otherwise, all protein concentrations were expressed as protomers. Hence, an assay containing 1 µM GroEL and 1 µM GroES, contained 71.4 nM of tetradecameric GroEL_14_ complexes, corresponding to 142.8 nM of active heptameric catalytic sites (GroEL_7_ cavities), which could be transiently covered by as many (142.8 nM) functional GroES_7_ heptamers during the ATP-driven catalytic cycle.

Chaperone stoichiometries in assays: GroEL and GroES protomers were always equimolar, as in bacteria and mitochondria^15^. For DnaK, DnaJ, GrpE and ClpB molar ratios were 5:1:1:2 Native reporter enzymes (MDH, CS, Luc) from stocks were diluted to the indicated final concentrations (0.25–2.5 µM) and incubated in thin-walled PCR tubes in a thermocycler at constant indicated temperatures, in the presence of the chaperones and ATP, typically for about two hours, but in the experiment in Fig.4, for up to 8 hours. At indicated times, aliquots were assayed in a spectrophotometer or a luminometer. Linear enzymatic rates were converted into molar concentrations (nM) of natively refolded chaperone products as explained below.

For experiments following the native refolding of pre-unfolded reporter enzymes, a 40-fold excess of MDH reporter enzyme was preincubated in 7 M urea and 20 mM DTT for 5 min at 25°C. Typically, 2 µl were then flush-diluted and mixed into 78 µl of pre-warmed refolding buffer containing chaperones and ATP, as indicated, and further incubated at the indicated temperatures. In contrast to Guanidium HCl that affects GroEL-GroES activity, the 175 mM traces of Urea were inconsequential for the chaperonin assay^78^.

### Calculation of the scale for graphs where enzymatic activity is reported

For a given total concentration [E] of the enzyme E in solution, (E being MDH, citrate synthase or luciferase), the maximal activity measured throughout the whole set of experiments represented in a figure is considered to represent [E], and all other activities are thus proportionally scaled when expressed as a concentration.

## Acknowledgments

We thank summer project students Patrick Inns and Ivanoé Koog for setting up the Citrate Synthase enzymatic assay and performing key preliminary and control chaperone assays. This project was financed in part by Swiss National Science Foundation Grants 140512/1, 31003A_156948 and by Grant C15.0042 from the Swiss State Secretariat for Education Research and Innovation to P.G. and B.F. Grant 200020_163042 from the Swiss National Science Foundation to P.D.L.R and A.S.S.A.B. acknowledges the support of the French Agence Nationale de la Recherche (ANR), under grant ANR-14-ACHN-0016.

## Supplementary Information for “Molecular chaperones inject energy from ATP hydrolysis into the non-equilibrium stabilisation of native proteins.”

Pierre Goloubinoff, Alberto S. Sassi, Bruno Fauvet, Alessandro Barducci, and Paolo De Los Rios

### The model

We define the following concentrations:

- [ND] is the concentration of native monomer bound into native (active) dimers;
- [NM] is the native monomer concentration;
- [I] is the less structured (unfolded) intermediate concentration;
- [M] is the misfolded monomer concentration;
- [A] is the concentration of proteins irreversibly trapped in aggregates;
- [CI] is the concentration of chaperone-intermediate complexes;
- [CM] is the concentration of chaperone-misfolded complexes;
- [C] is the concentration of free chaperones.

Then the model depicted in Fig.1 is defined by several equations describing the evolution of the various concentrations following various conformational and binding/unbinding transitions:

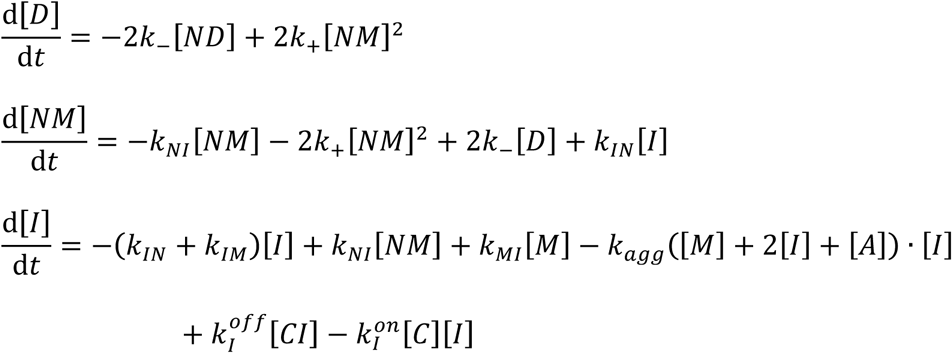

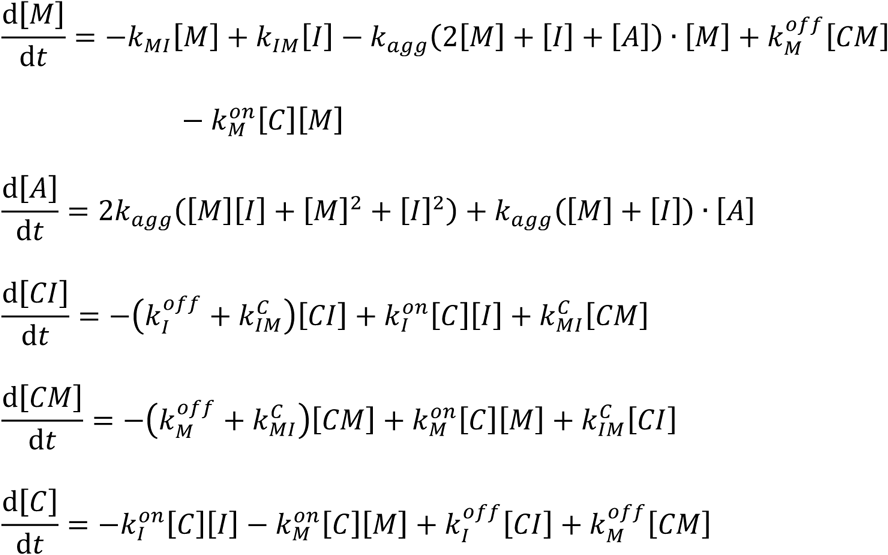

The rates of the reactions have been chosen so to reasonably reproduce the spontaneous denaturation of the substrate and the action of chaperones added at t=30’ in Fig.2A. The rates of the reversible chaperone-independent transitions are

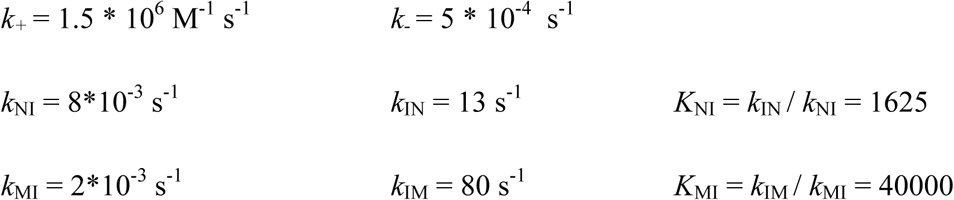

These values make dimers intrinsically much more stable than native monomers. Both native and misfolded monomers are much more stable than the (more) unfolded intermediates. Globally the non-native ensemble (misfolded and intermediates together) is more stable than the native ensemble (native monomers and dimers together).

Aggregation is a concentration-dependent process that proceeds from the non-native ensemble at a rate

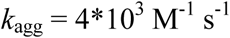

which compares with aggregation rates tabulated in the literature (see *e.g.* Powers *et al*.^1^). This rate is slow enough to account for non-native species to be recovered by non-disaggregating chaperones after several tens of minutes, but is fast enough to appear in the later stages of the spontaneous denaturation process.

Chaperone-mediated reactions deserve a special discussion. The rates of conversion between the misfolded and intermediate conformations, while bound to the chaperones, have been chosen as

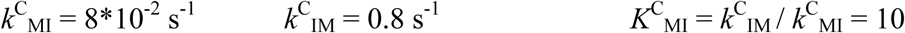

These rates imply a relative “unfolding” or “decompaction” action by the chaperone, which is not enough to tilt the balance in favor of the intermediate, less structured state *per se.* The balance is shifted toward the intermediate state with respect to the balance for substrates not bound to chaperones 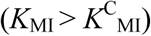.

In the model, non-equilibrium affects the dissociation rates of the chaperone/substrate complex. Indeed, when applying a fundamental rule that holds for all cycles leading to biochemical transformations, we obtain

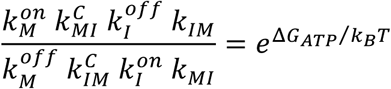

As a consequence, fixing the I-to-M and M-to-I transformation rates when in solution and when bound to the chaperone, and thus their equilibrium constants *K_MI_* and 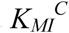, the substrate-chaperone association and dissociation rates are related by the relation

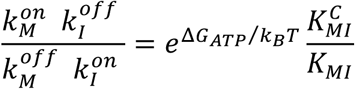

We chose identical chaperone association rates for the intermediate and misfolded conformations that are in line with the typical binding rates for protein complexes (10^6^ M^-1^ s^-1^):

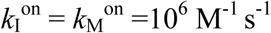

which implies that the injected energy ΔG_ATP_ modulates the relation between the unbinding rates:

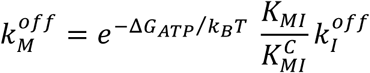

In the model, we chose:

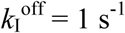

that in turn directly determines the value of 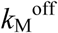 depending on ΔG_ATP_. ΔG_ATP_=0 at equilibrium; ΔG_ATP_=7.7 kcal/mol in Figs.3A–D, 4B and 5C.

Remarkably, the concerted hydrolysis of 7 ATP molecules by a GroEL_7_ ring liberates 63–84 kcal/mol ^2^. Yet, not all of it must be taken into account in the cycle: indeed, part of the energy is used to maintain also the internal conformations of the chaperone system (GroEL and GroES) away from their equilibrium concentrations^3^. Since those conformations are not accounted for in detail in the model, the energy they consume should in principle be subtracted from the one available to drive the cycle, ΔG_ATP_. Furthermore, as shown in Fig.5A, injecting more energy into the cycle would not produce significant differences that would be detectable from our experiments. Of course, it is important that the value of ΔG_ATP_ that we use does not exceed the one available from the hydrolysis of 7 ATP molecules.

In Fig.5A, ΔG_ATP_ is variable (as indicated on the horizontal axis).

In Fig.3E, ΔG_ATP_ depends on the concentrations of ATP and ADP, as indicated in the main text and in the figure legend (we kept the concentration of inorganic phosphate equal to the concentration of ADP, in the assumption that it was only produced by ATP hydrolysis), and we allowed the concentration of ATP and ADP to evolve in time. ATP was hydrolyzed (and thus its concentration decreased) at a rate proportional to the flux of dissociation of the chaperone-intermediate state complex:

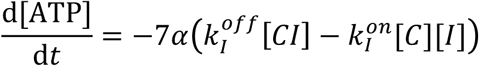

where we chose α=12 to best reproduce the results.

In the model, we neglected possible unproductive hydrolysis cycles: hydrolysis by unbound chaperones; binding of intermediate states and release of misfolded states; or binding of intermediate or misfolded states and their release after hydrolysis without causing any structural changes. Since each hydrolysis cycle consumes at least 7 ATP molecules per ring, the value α=12 implies that only 1 out of 12 hydrolysis cycles was productive. Once again, it is important to stress that our goal here is not a detailed accounting of all the rates. Rather our goal was to highlight the non-equilibrium nature of the action of chaperones, all while respecting the basic energy budget of the process.

### Meaning of the “apparent” free energy

According to equilibrium thermodynamics, the free energy difference between the native and non-native ensembles of proteins is given by

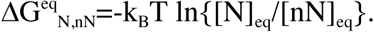

In a non-equilibrium stationary state the free energy, which is an equilibrium quantity, is not a rigorously defined. Nonetheless, we can use the same formula that is customarily used at equilibrium to define an “apparent” free energy difference. Indeed, an observer who was tasked with measuring this free-energy difference from the concentrations of the two ensembles, and who was unaware that there was an energy consuming process going on, would define the free-energy difference precisely according to the usual scheme using the above formula.

**Fig.S1.**
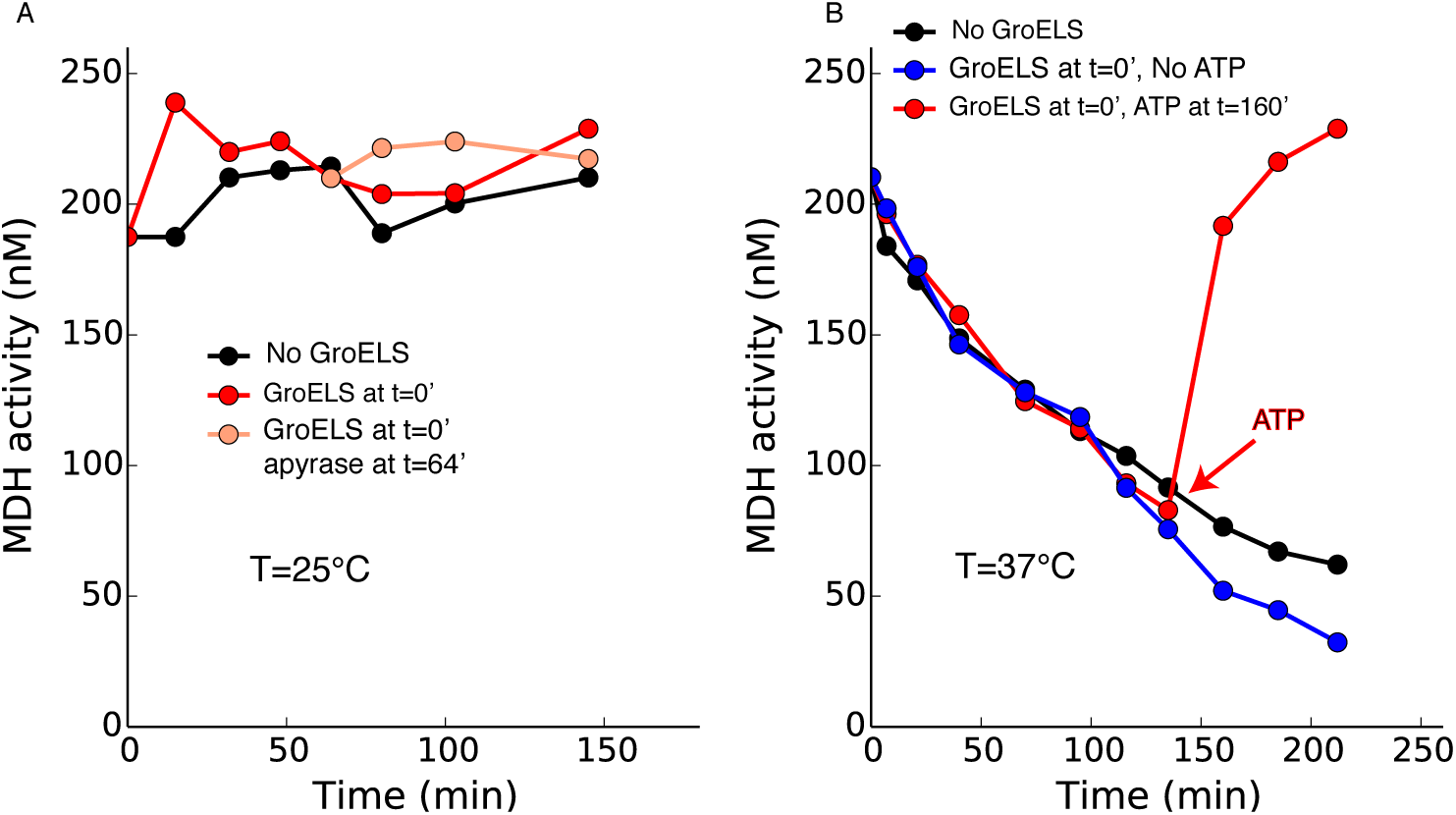
Behavior of MDH at 25°C and at 37°C. A) MDH (250nM) was stable at 25°C, both in the absence (black circles) and presence of GroELS and ATP (red circles), and of the addition of apyrase (pink circles). B) At 37°C, MDH (250 nM), was not stable in the absence of GroELS (black circles). In the presence of GroELS but without ATP, MDH lost activity roughly at the same rate as in the absence of chaperones (blue circles, and red circles up to t=160’). Addition of ATP at 160’ led to the recovery of almost all the lost activity (red circles).

**Fig.S2.**
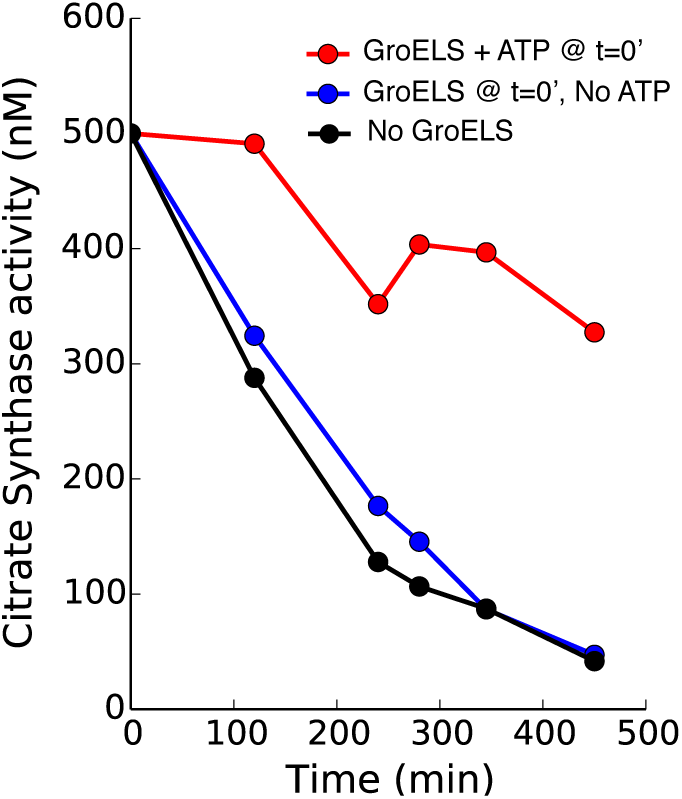
Behavior of Citrate Synthase at 44°C. Citrate Synthase (CS, 500 nM) spontaneously loses its activity at 44°C (black circles). The initial addition of GroEL (1 µM) and GroES (4 µM)^4^, but no ATP, did not change significantly the denaturation rate of CS (blue circles). The further initial addition of ATP and of a regeneration system slowed down the denaturation of the CS solution (red circles), consistently with our observations for MDH, thus strengthening our conclusions that GroELS can inject energy in the system, and transduce it in the form of an enhanced stability of the native state of its substrates.

**Fig.S3.**
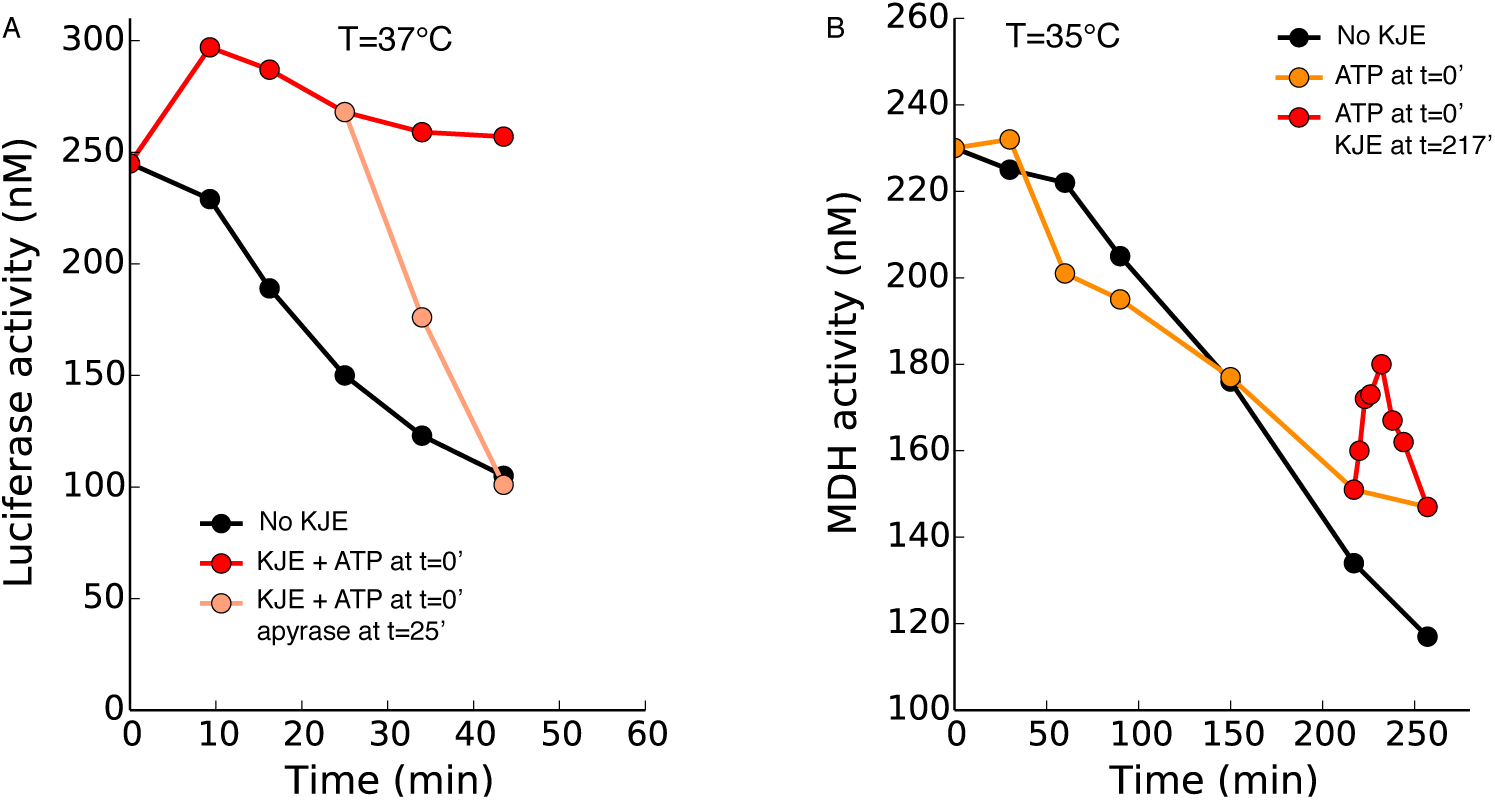
Reactivation and regeneration action of Hsp70 on Luciferase and MDH. A) Luciferase (0.3 µM) is intrinsically unstable at 37°C (black circles). The initial addition of DnaK, DnaJ and GrpE (2 µM, 0.4 µM and 0.4 µM respectively) and ATP regeneration system maintains Luciferase natively stable (red circles) until the later addition of apyrase leads to its denaturation (pink circles). Loss of activity in the presence of apyrase appears faster than in its absence because apyrase, rapidly hydrolyzing ATP, also affects the luciferase assay. B) MDH (0.25 µM) spontaneously loses activity at 35°C in the absence of chaperones (black circles). In the presence of ATP but no chaperones it also denatures spontaneously, whereas the addition of DnaK, DnaJ and GrpE (2.7 µM, 0.45 µM and 0.9 µM respectively) after 217’ produces a transient reactivation that, given the spontaneous tendency of MDH to denature, is only compatible with the injection of energy from ATP hydrolysis by KJE, albeit only transiently.

**Fig.S4.**
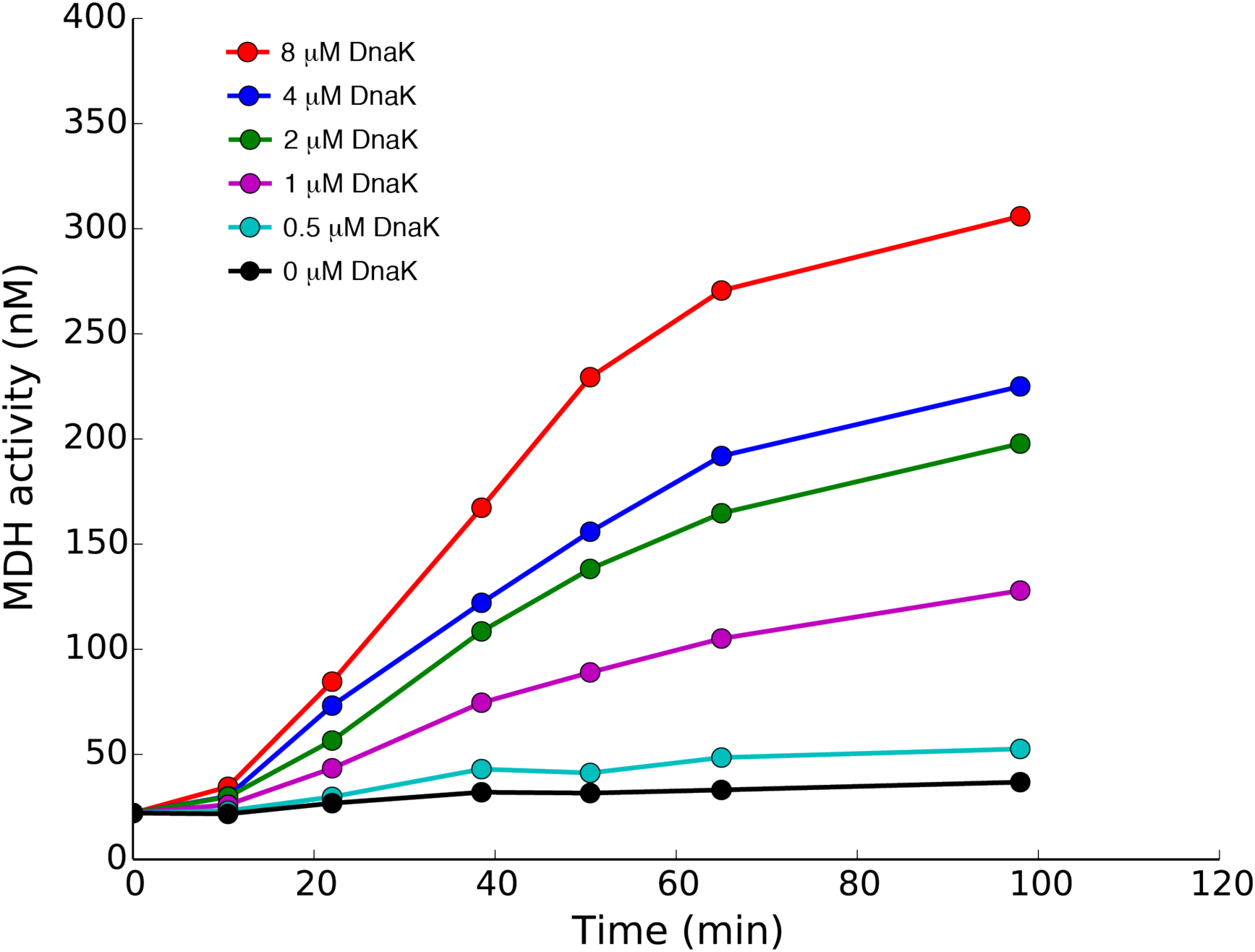
The full ClpB, DnaK, DnaJ, GrpE, GroELS network reactivates pre-aggregated MDH at the denaturing temperature 37°C. At non-native temperature, 37°C, pre-denatured and aggregated MDH (0.5 *µ*M) does not spontaneously reactivate in the absence of chaperones, nor in the presence of ClpB (2 *µ*M), GroEL and GroES (1 *µ*M each), DnaJ (0.8 *µ*M) and GrpE (0.8 *µ*M) but no DnaK (black circles). Increasing concentrations of DnaK allow the reactivation and maintenance of progressively more MDH, highlighting on the one hand the role of DnaK in the transduction of energy in the full chaperone network, and on the other hand showing that, when the full network is in place, it is powerful at injecting energy from ATP hydrolysis in the system and at turning it into an enhanced stabilization of native MDH, even in the presence of aggregates.

**Fig.S5.**
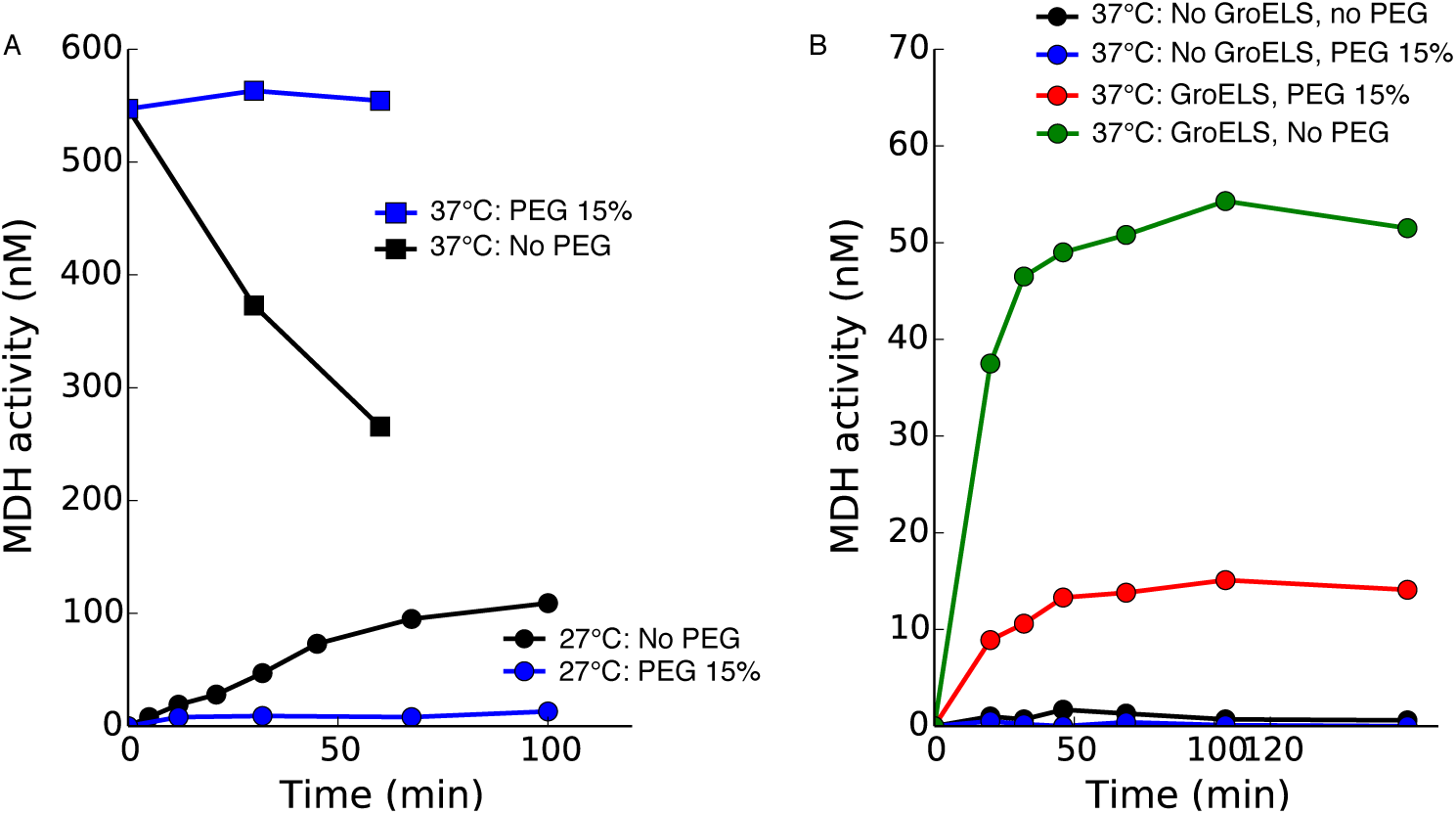
The effects of PEG on the stabilization of compact conformations highlight the role of GroELS. A) At 27°C, urea predenatured MDH (0.56 *µ*M) spontaneously refolded in the absence of PEG (black circles). In the presence of PEG 6000 (15%), spontaneous refolding was inhibited (blue circles). At 37°C, native MDH (0.56 *µ*M) denatured spontaneously in the absence of PEG (black squares). Addition of PEG 6000 (15%) inhibited denaturation (blue squares). The effects of PEG in this panel are consistent with the broadly accepted hypothesis that crowding agents like PEG destabilize unfolded, expanded conformations, thus increasing the transition times between compact conformation that must expand before converting into each other. B) At 37°C, urea predenatured MDH (0.56 *µ*M) does not spontaneously recover any activity (black circles), in keeping with its thermodynamic instability. The presence of PEG 6000 (15%), a crowding agent, does not help refolding, implying that PEG, by itself, does not stabilize the native state (blue circles). In the absence of PEG but in the presence of GroEL and GroES (3 *µ*M each) and ATP, some MDH activity can be readily recovered (green circles). Remarkably, PEG decreases the effectiveness of GroELS (red circles). Because according to the common view, PEG and other crowding agents favor compact conformations, these results suggest that the compaction action of GroELS counters the compaction action of PEG, providing support for the view that de-compaction is a crucial step, although possibly not the only one, in the mechanism of action of GroELS.

